# Laser particle barcoding for multi-pass high-dimensional flow cytometry

**DOI:** 10.1101/2022.06.03.494697

**Authors:** Sheldon J.J. Kwok, Sarah Forward, Marissa D. Fahlberg, Sean Cosgriff, Seung Hyung Lee, Geoffrey Abbott, Han Zhu, Nicolas H. Minasian, A. Sean Vote, Nicola Martino, Seok-Hyun Yun

## Abstract

Flow cytometry is a standard technology in life science and clinical laboratories used to characterize the phenotypes and functional status of cells, especially immune cells. Recent advances in immunology and immuno-oncology as well as drug and vaccine discovery have increased the demand to measure more parameters. However, the overlap of fluorophore emission spectra and one-time measurement nature of flow cytometry are major barriers to meeting the need. Here, we present multi-pass flow cytometry, in which cells are tracked and measured repeatedly through barcoding with infrared laser-emitting microparticles. We demonstrate the benefits of this approach on several pertinent assays with human peripheral blood mononuclear cells (PBMCs). First, we demonstrate unprecedented time-resolved flow characterization of T cells before and after stimulation. Second, we show 33-marker deep immunophenotyping of PBMCs, analyzing the same cells in 3 back-to-back cycles. This workflow allowed us to use only 10-13 fluorophores in each cycle, significantly reducing spectral spillover and simplifying panel design. Our results open a new avenue in multi-dimensional single-cell analysis based on optical barcoding of individual cells.

## Introduction

Fluorescence-based flow cytometry has been a workhorse in the single-cell analysis of surface markers, intracellular cytokines, intranuclear proteins such as transcription factors, and cell cycles. Continuing advances in high-speed fluidics and multi-color optics, as well as fluorophore chemistry, has enabled high-parameter measurement (up to ~30 markers) at high speed (>10,000 cells per second) and low cost. While these are major advantages in throughput and cost^1,2^ over technologies such as single-cell mass cytometry and sequencing-based proteomic analysis, flow cytometry is facing significant challenges in meeting the growing demand to measure more protein markers per cell. Highly multiplexed measurements of immune cells to characterize dozens of different cell types have proven to be critical in the development of immunotherapies and vaccines^3–6^, as well as detection of minimal residual disease in leukemia^7^. However, high-marker analysis (>30 protein markers) is challenging due to the ambiguity caused by spectral spillover between fluorophores, often requiring months-long optimization of fluorophore-antibody combinations and instrument settings^8,9^. Clinical laboratories that have the labor and time available to optimize a high-marker panel may still lack the expertise to design and select the appropriate reagents. Limited availability of well-validated fluorophores (colors) is an additional barrier, especially for clinical applications which require the use of reagents that meet FDA manufacturing requirements. For these reasons, most clinical laboratories use standardized panels for immunophenotyping that typically range from 4-12 colors, which restricts the types of cells that can be detected at once and increases the number of cell samples required^10–12^.

A major advantage of flow cytometry is that, unlike mass cytometry and sequencing, cells are not destroyed during optical acquisition. Flow sorters rely on this nondestructive feature. However, current flow cytometer *analyzers* are typically used for one-time measurement of cells. Measuring cells twice using a flow sorter is in principle possible, but single-cell information would be lost in the cell collection process. This limitation, which has not been openly recognized, makes current flow cytometry unable to address the ever growing need to acquire high-dimensional data and temporal responses of single cells^13–15^.

Here, we introduce a drastically new approach in flow cytometry based on optical barcoding of individual cells. Barcoding techniques have been used previously in cytometry for tracking different samples, enabling pooling of samples for faster analysis. These techniques, relying on fluorescence intensity differences in flow cytometry^16^ or a limited set of radioisotopes in mass cytometry^17^ are only suitable for tracking tens of samples at a time. Here, we use laser particles (LPs), recently developed laser-emitting microparticles^18^, to tag and track up to millions of cells at a time. This approach enables multi-pass flow cytometry, in which the same cells are measured multiple times, using each cell’s unique optical barcodes to align and concatenate data from different measurements. We use this method to acquire flow cytometry data from human peripheral blood mononuclear cells (PBMCs). First, we apply our multi-pass approach to a common assay involving *in vitro* stimulation of T-cells in PBMCs. We demonstrate unprecedented characterization of the same T cells before and after stimulation, enabling quantification of marker downregulation. Second, we applied multi-pass cytometry to high-marker analysis. We developed a broad immunophenotyping panel optimized for 3 measurement cycles of leukocyte populations. Our “cyclic” approach greatly simplifies high-parameter analysis by requiring a far fewer number of fluorophores for the same number of markers. We performed this 33-marker assay on live human PBMCs from a healthy donor and validated our results against published data.

## Results

### Multi-pass flow cytometer instrumentation

Figure 1 illustrates the general workflow of multi-pass flow cytometry along with the optical measurements of cellular barcodes and fluorescent reagents. First, cells are mixed with excess LPs in solution to label each cell with a unique, random combination of LPs. Next, cells are stained with a first set of antibody-fluorophores and then loaded into a flow cytometer capable of exciting and detecting the laser emission from LPs and, also, collecting the cells after the flow measurement. We have built such a multi-pass flow cytometer using a near-infrared pump laser (1064 nm) to stimulate the laser emission of LPs and four fluorescence excitation lasers (405 nm, 488 nm, 560 nm, and 630 nm) to elicit fluorescence. Cells flow across the laser beams in a hydrodynamically focused stream at a velocity of ~3.4 m/s. The fluorescence signal is split by dichroic filters and detected by avalanche photodiodes, while the lasing signal is detected by an infrared linescan spectrometer. Following data acquisition, the antibody-fluorophores in the collected cells are deactivated by either the photobleaching of the fluorophores or the release of the antibodies from the markers. For the next cycle of measurement, the cells are stained with a subsequent set of antibody-fluorophores and loaded back into the flow cytometer. In the following sections, we describe each of the major workflow steps in detail.

**Fig. 1.**
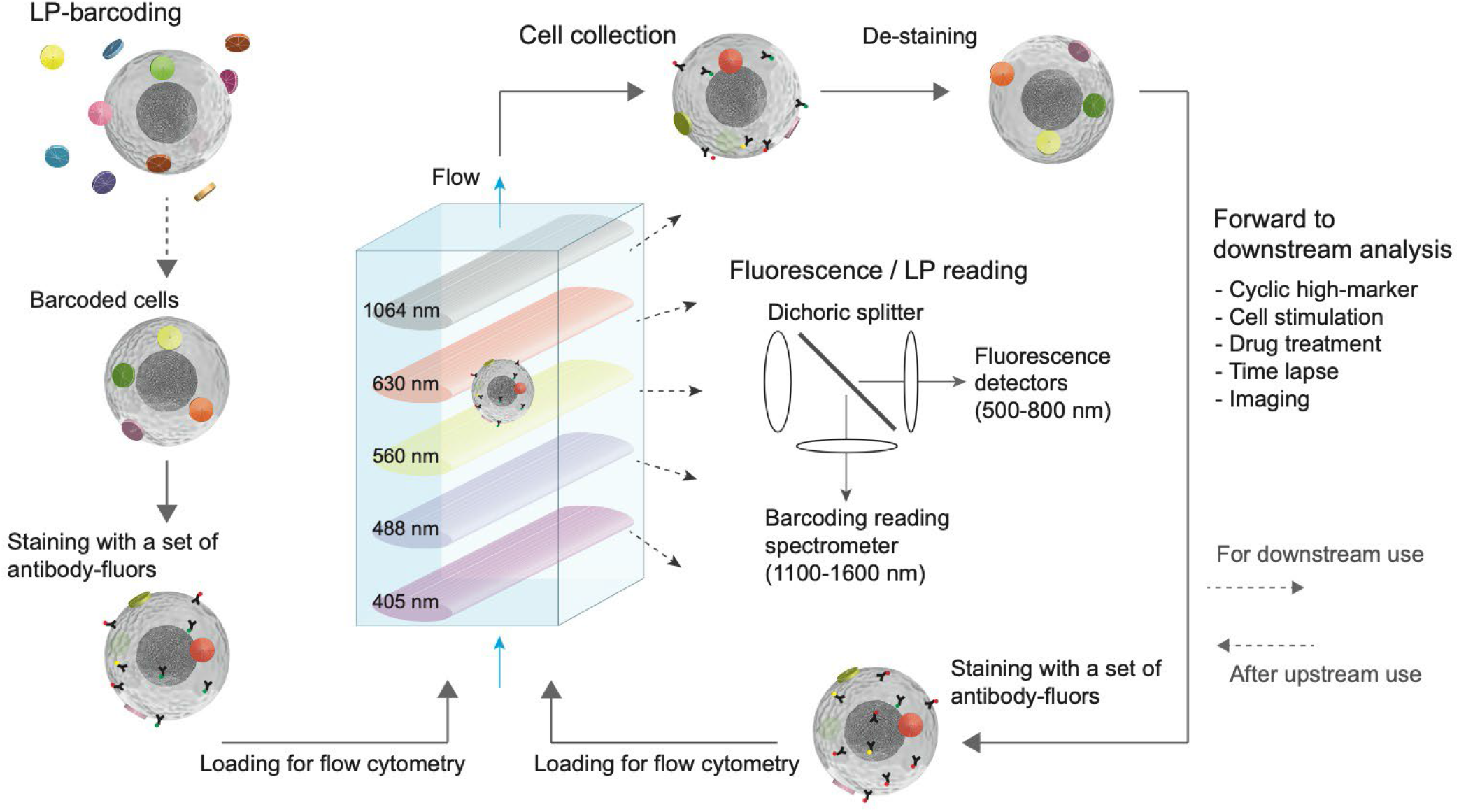
Schematic of multi-pass flow cytometry. The major steps include: tagging cells with laser particles (LPs) to yield LP-barcoded cells which typically have 3+ LPs, then staining cells with a first set of fluorophore-conjugated antibodies, loading the barcoded and stained cells into a flow cytometer that detects fluorescence signals using multiple excitation lasers and LP lasing signals using a pump laser and spectrometer, then collecting the cells, de-staining (e.g. photobleaching of the fluorophores or chemical release of antibodies), re-staining the cells with another set of antibodies and measuring them again in the flow cytometer. In addition to flow cytometry, this technology can be expanded to further downstream and upstream analysis of the barcoded cells.

### Laser-particle tagging of live human PBMCs

As LPs, we used InGaAsP microdisks^18^ of 1.6-1.9 μm in diameter and 220-290 nm in thickness (Fig. 2a). Six different compositions of bulk InxGa1-xAsyP1-y epitaxial layers were used to ensure each LP emits a lasing peak between 1150 to 1550 nm (Fig. 2b), which leaves the entire visible and near-infrared wavelength ranges free for fluorescence labeling. We first coated the semiconductor microdisks with a ~50 nm layer of SiO2 to ensure stability and confer biocompatibility. We have previously shown that silica-coated LPs can be internalized into a variety of cell types with overnight incubation^18^. To shorten the tagging time, we functionalized the silica coating surface with polyethylenimine (PEI), a cationic polymer known to bind to cell membranes and used for transfection^19,20^. We found that PEI-silica coated LPs were efficiently attached to live human PBMCs within 1 h of incubation (Fig. 2a). A given cell is defined as barcoded if it is tagged with 3 or more LPs. Typically, around 85% of cells were tagged with 1 or more LPs, and around 50% tagged with 3 or more LPs (Fig. 2c). Another approach for LP tagging involves antibody-based targeting of LPs to the cell surface through biotin-streptavidin coupling (Supplementary Fig. 1). PEI-based tagging was the primary method used in this study.

**Fig. 2.**
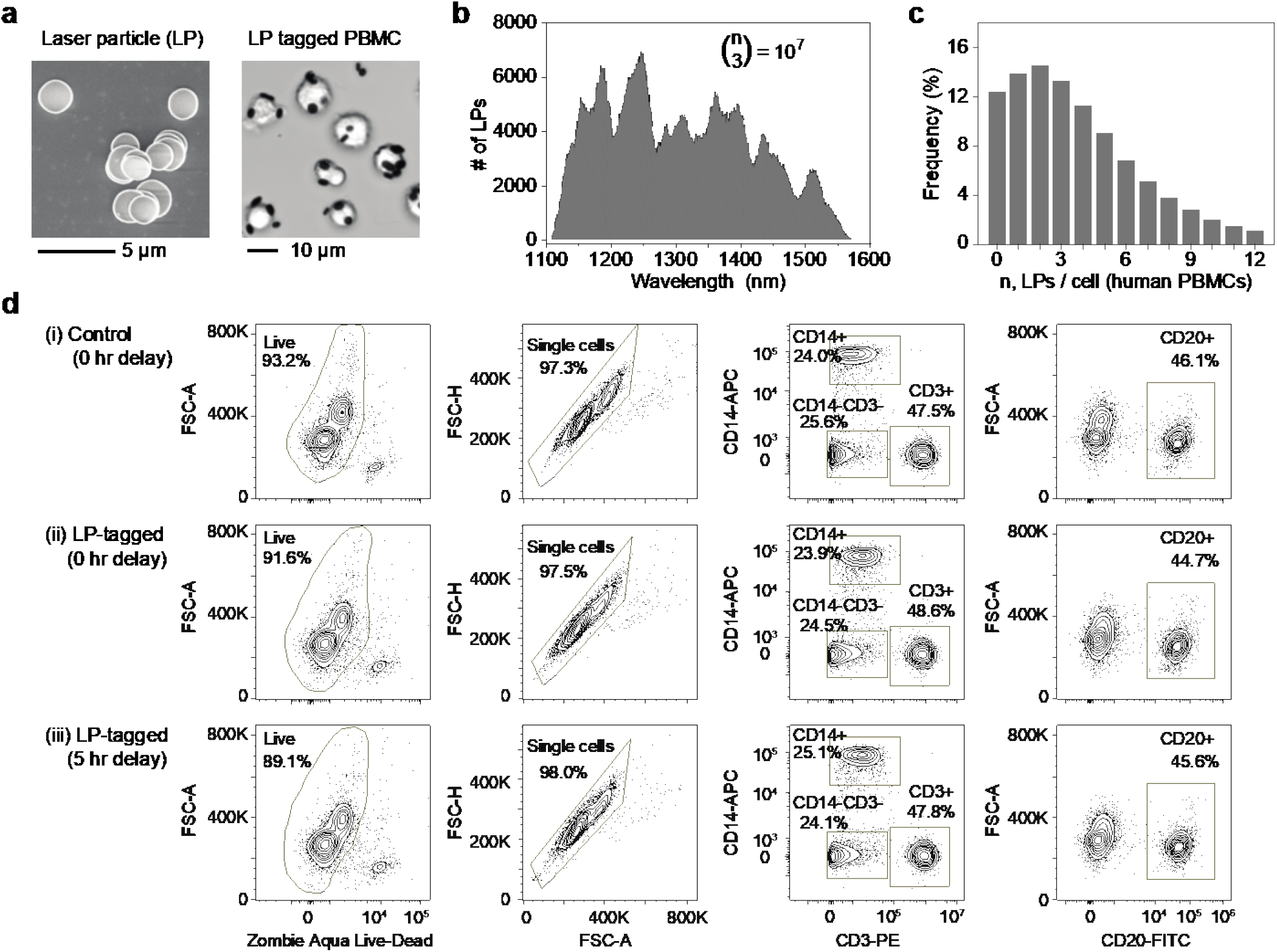
Tagging of Human PBMCs with LPs. (a) Left: Scanning electron micrograph of polymer-silica-coated LPs. Right: Optical image of live human PBMCs tagged with LPs. (b) Lasing wavelength distribution of the LPs comprising 10^7^ distinguishable barcodes when used in combinations of 3 or more. (c) Histogram showing distribution of LPs / cell, with over 50% of cells having 3 or more LPs. (d) Immunophenotyping data comparing cell viability, singlet purity, and frequency of monocytes, T cells and B cells for (i) control, untagged cells at 0 h, (ii) LP-tagged cells at 0 h, and (iii) LP-tagged cells at 5 h post tagging.

Using a viability dye that assesses membrane integrity, we found that cell viability is nearly unchanged from 93.2% before to 91.6% after LP tagging. Storing the LP-tagged samples in standard wash buffer at 4°C for five h reduces viability slightly further to 89.1% (Fig. 2d). There was no measurable difference in singlet purity between control and LP-tagged samples at 0 and 5 h, indicating LP tagging does not increase cell-cell aggregation. To test whether LP-tagging affects cellular phenotypes, we also compared the expression of major immune markers including CD45 (pan immune marker), CD14 (monocyte marker), CD3 (T-cell marker) and CD20 (B cell marker). There were no significant differences in the population percentages of any of these markers when comparing control and LP-tagged samples at 0 and 5 h (Fig. 2d). Furthermore, using CD3-KromeOrange, we found that the median fluorescence intensity decreased approximately ~5% for cells tagged with 3-5 LPs, and ~15% for cells tagged with 10 LPs (Supplementary Fig. 2).

### Repeated identification of the same cells

Cells are collected after each measurement so that the same cells can be measured again in the subsequent cycle. In conventional flow cytometers employing hydrodynamic focusing, cells flow through a glass flow cell along with sheath fluid and are then diverted into waste following analysis. We developed a novel, cell-collecting fluidic channel that recovers all the cells in the focused core stream of 10-20 μm width at the exit of the flow cell (Fig. 3a-b). The collection channel consists of a 127 μm diameter needle, followed by a polypropylene-based flexible tube connected to a peristaltic pump that controls the flow rate of the collected stream (Fig. 3c). With sample input flow rates of 30 μL/min, a sheath flow rate of 9 mL/min, and a collection flow rate of ~400 μL/min, we were able to collect nearly 100% of the input cells into a tube. To ensure viability of cells in the tube during flow acquisition, we used phosphate-buffered saline (PBS) as the sheath fluid, and the cells were collected in serum-supplemented buffer. Immediately after acquisition, the cells are washed and resuspended in standard flow cytometry staining buffer. Overall, the cell collection and washing process typically recovers 95% ± 2% of live human PBMCs (Fig. 3d). There was no significant change to the viability of human PBMCs after 1, 2 or 3 cycles of cell capture (Supplementary Fig. 3).

**Fig. 3.**
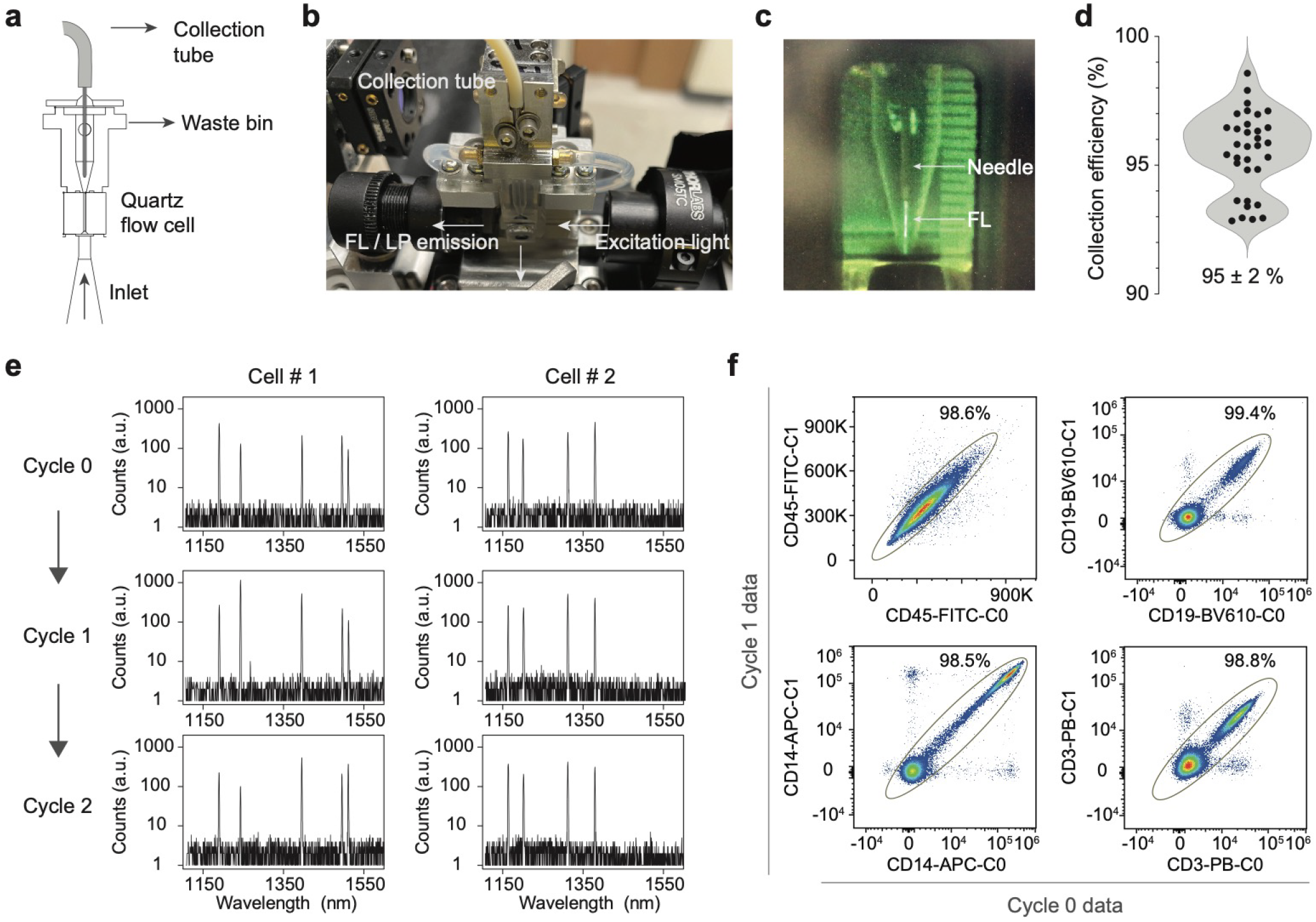
Repeated measurements of the same cells. (a) Schematic and (b) image of modified fluidics to incorporate a needle and collection tube for cell capture. (c) shows fluorescein dye flowing into the needle. (d) Cell collection efficiency of live CD45+ human PBMCs over 32 trials of capturing cells using the modified flow cell, with mean of 95 and standard deviation of 2%. (e) Representative lasing spectra of two LP-tagged cells measured repeatedly over 3 cycles. An algorithm was used to match cells between measurements using the lasing wavelengths. (f) Validation of matching LP spectral barcodes. LP-tagged cells stained with antibody-fluorophores were measured in successive cycles (Cycle 0 and Cycle 1) and then matched using their lasing spectra. Plots show strong correlation between fluorescence signals measured in the two cycles.

Cells measured in different passes were matched using their LP barcodes. Our spectrometer measures the lasing spectra with resolution of ~ 0.5 nm. Given that single LPs provide about 800 distinguishable lasing peaks (from 1150 to 1550 nm), a combination of three random LPs per cell in principle can provide 800C_3_ = 8.5×10^7^ unique spectral barcodes, sufficient for tracking a population of 1,000,000 cells with < 2% of duplication-induced error (loss). LP barcodes measured in different passes were matched by extracting the peak wavelengths and computing the probability of a match by comparing pairs of measured spectra. Each potential match is scored depending on the lasing wavelengths detected and their emission amplitudes (see Methods). Examples of matched spectra over three cycles are given in Fig. 3e. We validated our approach by staining LP-tagged PBMCs with major immune markers (CD45, CD14, CD3 and CD19), measuring the same cells twice using the modified flow cytometer, and comparing the fluorescence data of cells that were matched using the LP barcodes. We defined an apparently correctly matched population in which the fluorescence intensities of the same cells measured in Cycle 0 and Cycle 1 are strongly correlated (Fig. 3f). This population ranged from 98.5% to 99.4% of the cells. The remaining 0.6-1.5% of cells that were apparently correctly matched appear as noise and do not meaningfully affect data quality and resolution. We further validated our approach by verifying whether LP barcodes could be used to keep track of sample identity. Data from three separately acquired samples was pooled together and matched. Less than 2% of the matched cells were erroneously identified as belonging to two different samples (Supplementary Fig. 4). Our validation data shows our approach can track and match cells between different measurements with high accuracy.

### Time-resolved measurements of T cell activation

One potentially impactful application of multi-pass flow cytometry is time-lapse flow analysis. Multi-pass flow cytometry can be used to measure changes in marker expression of individual cells between subsequent cycles due to various biological processes either naturally occurring or artificially induced, such as cell division, incubation, drug treatments, cell-cell interactions, or stimulations. To measure the same markers in successive measurements, we employed chemically releasable antibodies^21^ (Miltenyi Biotec, REAlease), which are designed to be released from live cells with the addition of a chemical reagent.

We explored time-resolved flow cytometry analysis of T cells upon cell stimulation. Drug treatments or immunotherapies can alter the expression of protein markers on certain cell types, reflecting changes in activation state, viability, drug response or resistance^22–25^. While conventional flow cytometry can compare population differences between treated and untreated cells, it cannot identify changes to each individual cell, which is especially important for heterogeneous cell samples. In addition, changes in marker expression can prevent or impair identification of cell type post stimulation^26^. Using a basic T-cell panel, we measured two different samples of T-cells in human PBMCs from the same donor with and without stimulation with phorbol myristate acetate (PMA) and ionomycin. As shown in Fig. 4a, while CD4+CD3+ T cells can readily be identified prior to stimulation, CD4 expression is significantly reduced post stimulation, making it difficult to re-identify these cells. Furthermore, it is difficult with conventional flow cytometry to distinguish whether changes in marker are phenotypic switches or due to expansion or death of specific cell types.

**Figure 4.**
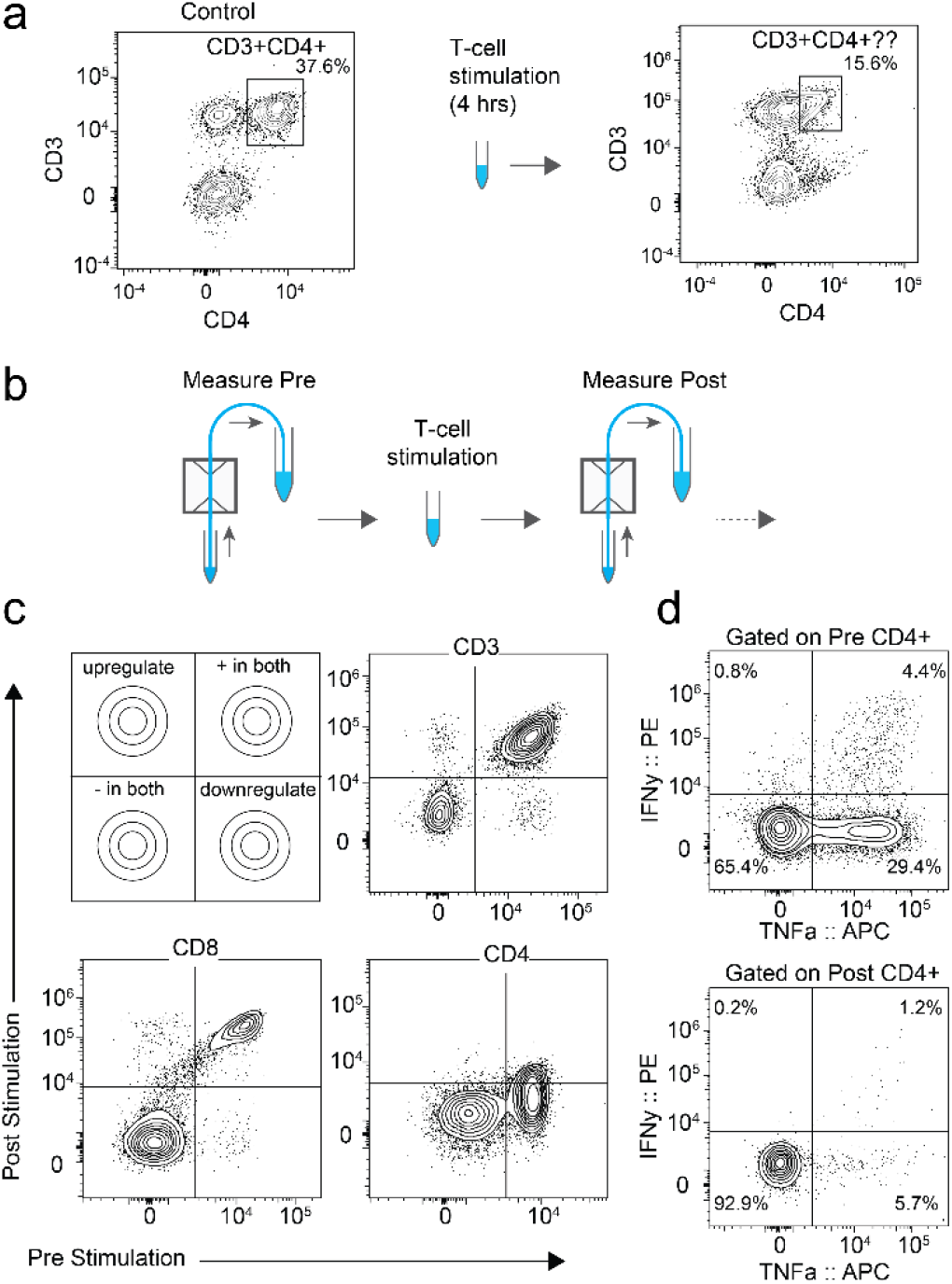
Time-resolved flow cytometry. (a) Conventional flow cytometry can identify CD3+CD4+ cells pre-stimulation (left), but T-cell stimulation results in loss of CD4 signal, making it difficult to identify CD3+CD4+ cells post stimulation (right). (b) Time-lapse flow cytometry in which T-cells are measured pre and post stimulation. (c) Plotting marker signals measured pre and post stimulation enable identification of markers that are upregulated or downregulated (top left). Plotting CD3 Pre vs CD3 Post revealed minimal change with stimulation (top right). Plotting CD8 pre vs. CD8 post also revealed minimal change with stimulation (bottom left). Plotting CD4 pre vs. CD4 post revealed significant downregulation (bottom right). (d) Gating on pre-stimulation CD4+ cells enables identification of cytokine-secreting cells (top). Gating on post-stimulation CD4+ cells identifies significantly fewer cytokine-secreting cells due to CD4 downregulation (bottom).

For time-resolved characterization of T-cell stimulation, we tagged human PBMCs with LPs, stained with chemically releasable antibodies (CD3, CD8 and CD4) and analyzed for baseline phenotyping (pre stimulation) (Fig. 4b). After the acquisition, the antibodies were removed, and the cells were stimulated for 4 h with PMA/ionomycin. The cells were then re-stained for the same surface markers in addition to intracellular cytokines, followed by a second-pass analysis with our cytometer post stimulation. Figure 4c shows expression of different markers (CD3, CD8 and CD4) pre and post stimulation for each cell. While CD8 and CD3 expression did not change significantly, we identified loss of CD4 expression on T cells that was easily identifiable during the first pass. Our time-resolved approach enables gating on pre-stimulation markers for downstream analysis, such as identification of cytokine-secreting cells that were CD4+ prior to stimulation (Fig. 4d).

### Photobleaching of common fluorophores

To measure different markers in each pass, antibodies need to either be removed or fluorescence signals need to be inactivated after each measurement so the cells can be re-stained with a different set of fluorophore-conjugated antibodies. While chemically releasable antibodies (Miltenyi Biotec, REAlease) are suitable, the limited portfolio of antibodies commercially available makes it difficult to use for high-marker panels. There are also a number of approaches for antibody stripping and/or iterative staining on fixed cells^27–30^ (CODEX, CyCIF, IBEX and 4i), but fewer well-validated options exist for live cells.

We developed and optimized an in-solution photobleaching method that was compatible with live cells. To minimize viability loss during photobleaching, we built a device that illuminates cell samples while actively cooling to near 4°C (Supplementary Fig. 5). Cells are suspended in a buffer containing an antioxidant that prevents reactive oxygen species from damaging cells (see Methods). Using broadband light emitting diodes (LED, 440-660 nm), we were able to photobleach a number of commonly used antibody-conjugated fluorophores (anti-CD45) in 3 to 25 minutes. A violet LED (400-420 nm) was needed to efficiently bleach violet-excitable fluorophores (Fig. 4a). After photobleaching, the fluorescence signal in the relevant channels (e.g., both donor and acceptor components for tandem fluorophores such as PE-Cy7) is similar to an unstained sample (Fig. 4b). Using 10 antibody-conjugated fluorophores at a time, the cell viability dropped slightly from 97.2% to 93.4% after a single bleach, and slightly further to 91.1% after two bleaches (Supplementary Fig. 6). We found that fluorophores conjugated to markers with higher antigen density generally tended to bleach more slowly, presumably because of limited local oxygen supply for bleaching. To verify that photobleaching does not change the relative expression of markers on cells and does not cause heterogenous cell loss, we performed immunophenotyping of live human PBMC sample after photobleaching and compared to a control. We found no significant differences between the percentages of CD4+ T cells, CD56+ NK cells, CD20+ B cells and CD14+ Monocytes (Fig. 4c).

### 33-marker, 3-cycle measurement of live human PBMCs

To demonstrate the utility of multi-pass flow cytometry for high-parameter flow analysis, we designed a 3-cycle, 33-marker deep immunophenotyping panel of human PBMCs using 10-13 fluorophores per cycle, with photobleaching between measurements (Fig. 5d). The 33 markers were chosen to enable identification of a variety of cell types including CD4+ T, CD8+ T, regulatory T, and γδ T, B cells, plasmablasts, NKT-like cells, NK cells, monocytes, innate lymphoid cells, and dendritic cells. For each cell type, differentiation and activation markers were included for sub-categorization, such as for naïve, memory, and effector T cells. Live human PBMCs from a healthy donor were used to acquire the data. The complete “cyclic” workflow included LP tagging, staining with Cycle 0, acquisition, photobleaching, re-staining with Cycle 1, acquisition, photobleaching, re-staining with Cycle 2 and a final acquisition. In this study, about 50% of the barcoded cells that are acquired in Cycle 2 were successfully matched with previous cycles.

**Fig. 5.**
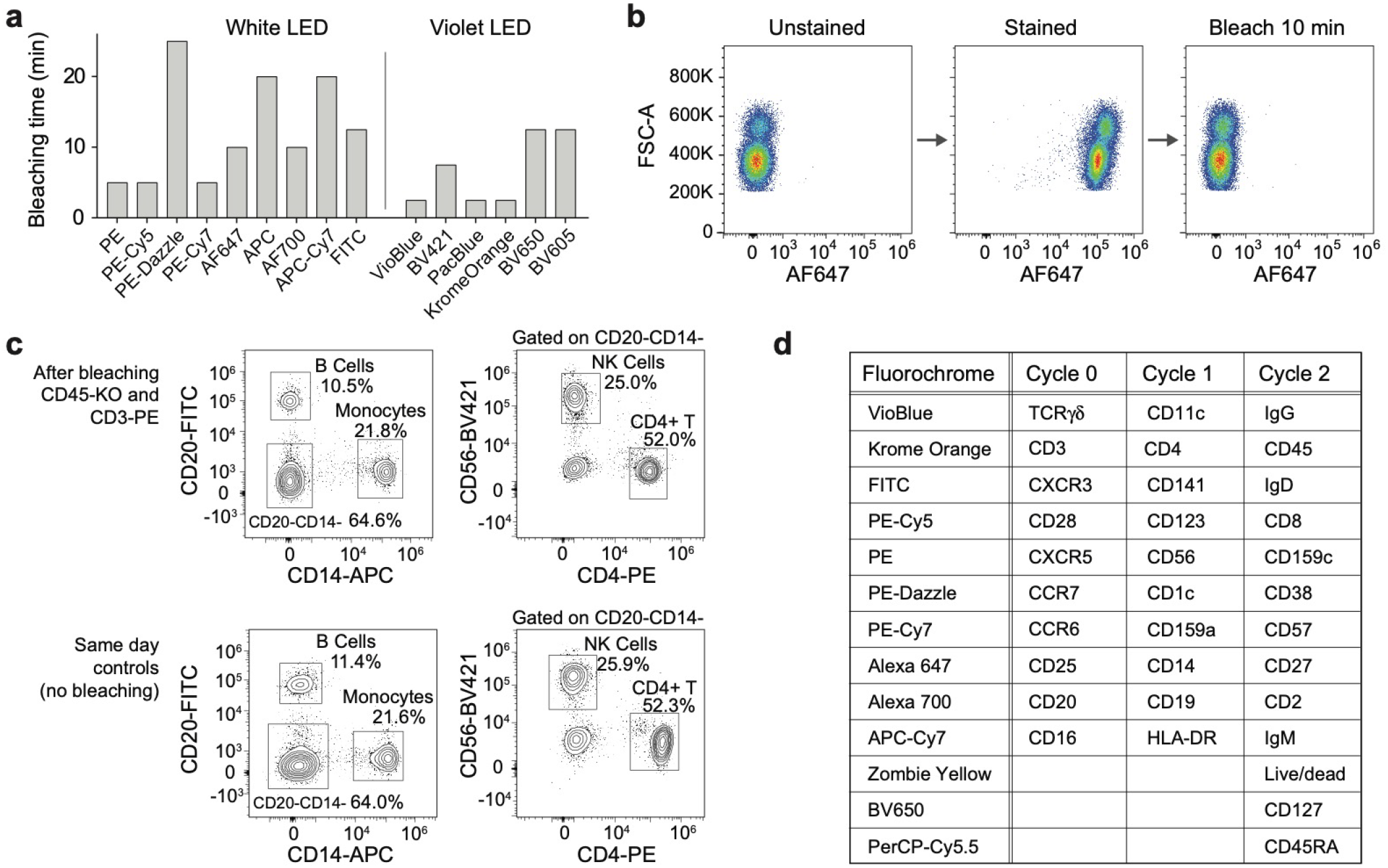
Fluorophore photobleaching for high-marker multi-pass cytometry. (a) Photobleaching time for fluorophores conjugated to CD45 excited by a broad-spectrum white light-emitting diode or a 405 nm light-emitting diode. (b) Representative data showing complete fluorescence signature erasure via photobleaching. Cells stained with CD45-AF647 photobleach in 10 mins to a median fluorescence intensity equal to that of an unstained cell sample. (c) Effect of photobleaching on live cells by re-challenging photobleached cells with antibodies targeting co-expressed markers. Cells stained with CD45-KrO and CD3-PE were fully bleached, re-stained with CD14, CD20, CD4 and CD56, and compared to a unbleached control sample. No significant differences were observed. (d) Panel design of high-marker experiment. Antibodies and fluorophores used in each cycle are shown.

Figure 6a visualizes the matched 33-marker data after dimension reduction using uniform manifold approximation and projection (UMAP). At least 6 distinct islands corresponding to different cell types were observed, with good separation consistent with high data quality. Cross-referencing the UMAP pattern with each measured marker yielded the expected major cell subsets in each island, including CD4+ T cells, CD8+ T cells, CD14+ monocytes, CD11c+ dendritic cells, CD123+ dendritic cells CD20+ B cells, and CD56+ NK cells. Repeating the 3-cycle measurement on different days with PBMCs from the same donor yielded similar results (Fig. 6b). To assess whether the cyclic workflow had any effect on the final results, we swapped the order of 10 antibodies between Cycle 1 and Cycle 2 on Day 3 (Fig. 6b). There were no significant differences in either the number of cell subsets identified or the quality of the data. The live cell fraction measured in Cycle 2 was found to be 87%. There were no significant differences in the viability in Cycle 2 for major cell types (Supplementary Fig. 7).

**Figure 6.**
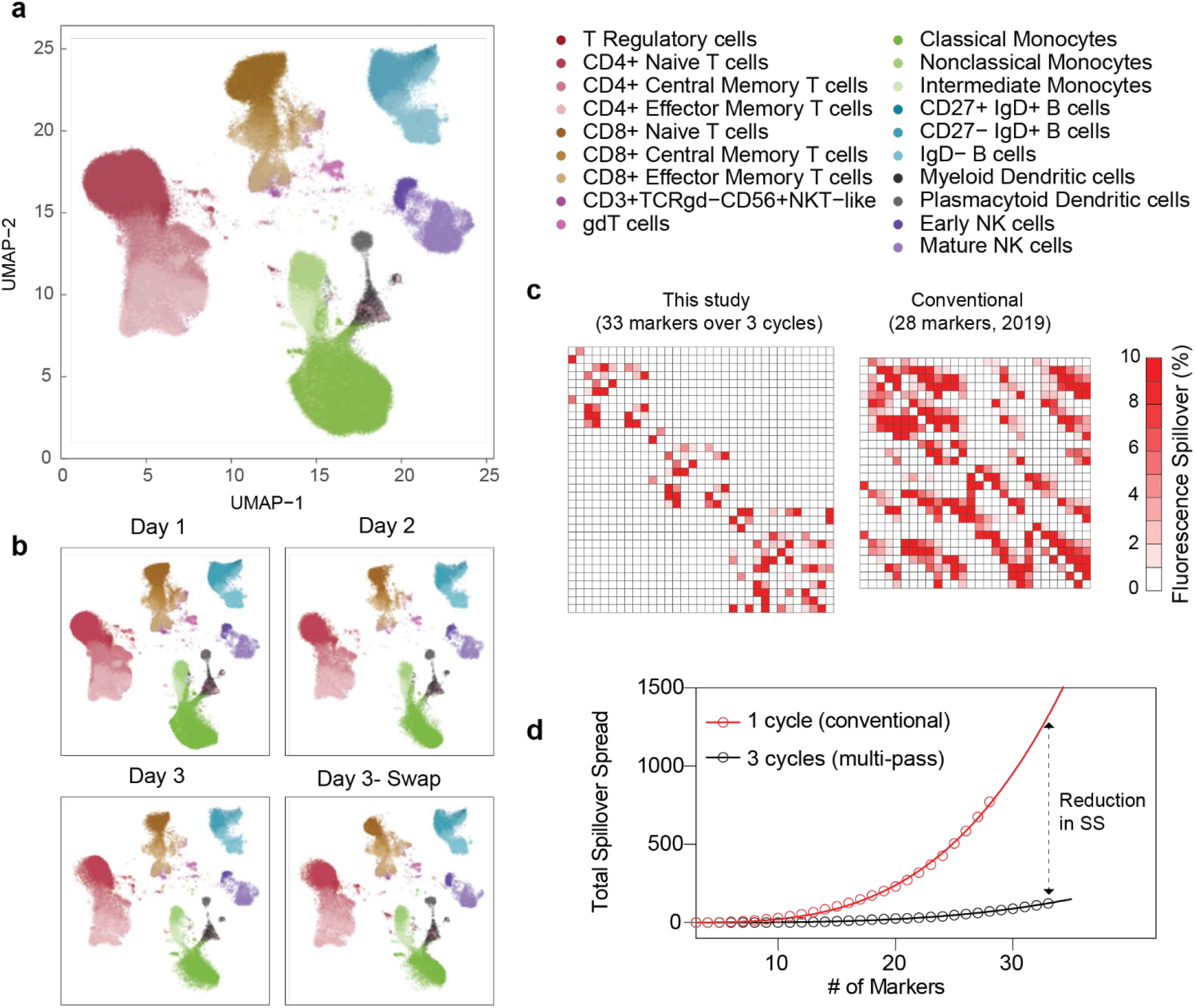
33-marker deep immunophenotyping of human PBMC over 3 cycles. (a) UMAP representation of data after being combined, matched, and cleaned using doublet discrimination, live/dead gating, and tight gating. Cell populations were manually gated and displayed on the UMAP by color. (b) UMAP representation of data taken on three different days (Day 1, Day 2, and Day 3) or after swapping reagents between different cycles (Day 3-Swap). No significant differences were observed with these datasets. (c) Compensation matrix of 33 marker, 3 cycle panel used in this study compared to conventional 28-marker panel in OMIP-060, ref 23. (d) Simulated total spillover spread computed for a high-marker panel using a multi-pass 3 cycle vs. a conventional 1 cycle workflow.

We computed the compensation matrix used in our study (33 markers over 3 cycles) and compared it to a 28-marker panel optimized for conventional single-pass flow cytometry^31^,^32^. As expected, there was significant reduction in spillover resulting from acquiring fewer colors over multiple passes (Fig. 6c). There were 136/1056 (12.9%) pairs with spillover greater than 0.5% in our panel vs. 393/756 (52%) pairs in the conventional panel. When comparing panel performance within a given instrument, spillover spread (SS) is typically used to indicate how well co-expressing markers can be resolved when stained with a specific combination of colors^33^. We found that the total amount of SS in high-parameter panels increases nonlinearly (power law with an exponent of about 3) with each additional color used (Supplementary Note 1). In contrast, a 3-cycle workflow had significantly reduced SS, over one order of magnitude times lower (Fig. 6d; Supplementary Note 1). Supplementary Figure 8 show the SS matrices of the 33 marker, 3-cycle panel from this study and the published 28 color panel^31^.

Figure 7 show the acquired scatter plots and our gating tree. A manual gating strategy was used to distinguish T, B, NK, and myeloid cell subsets, following guidelines from previously published datasets^34–38^. Briefly, T-cells and their memory subtypes were identified by surface expression of CD45RA, CCR7, CD27, and CD28 on CD3+CD4+ cells or CD3+CD8+ cells. CD4+ helper T cell subtypes were further differentiated by expression of CXCR3, CXCR5, and CCR6, and T regulatory cells were defined as CD127^lo^CD25+. We characterized unconventional T cells by expression of the γδTCR or CD56 on CD3+ cells. Monocytes were defined as CD3-CD19-CD20-CD56-HLA-DR+ cells that expressed CD14 and/or CD16, while dendritic cells were gated with the same lineage but were here defined as lacking expression of CD14 and/or CD16. They were further characterized by expression of CD123 (pDCs) or CD1c (mDCs). Of note, this strategy may miss small subsets of dendritic cells which co-express CD16. B cells were initially defined by expression of HLA-DR, CD19, and CD20, and lack of CD3 and CD56. We used expression of IgD, IgM, and IgG to differentiate B cells producing antibodies of different isotypes. Plasmablasts were identified by expression of CD19, lack of CD20 and co-expression of CD27 and CD38. Using a combination of CD16, CD56, NKG2A, and NKG2C, we identified early, mature, and memory NK cells. Our results are consistent with previously published findings on T cells^35^, B Cells^36^, Treg and myeloid cells^37^, as well as NK cells^38^. Importantly, we validated expected co-expression of markers by staining them in different cycles. For example, >95% of naïve CD4+ and CD8+ T cells that we have defined by CD45RA (Cycle 2) and CCR7 (Cycle 0) also expressed CD27 (Cycle 2) and CD28 (Cycle 0), as anticipated. In addition, all CD20+ (Cycle 0) B cells co-expressed CD19 (Cycle 1), consistent with expected healthy human phenotypes.

**Figure 7.**
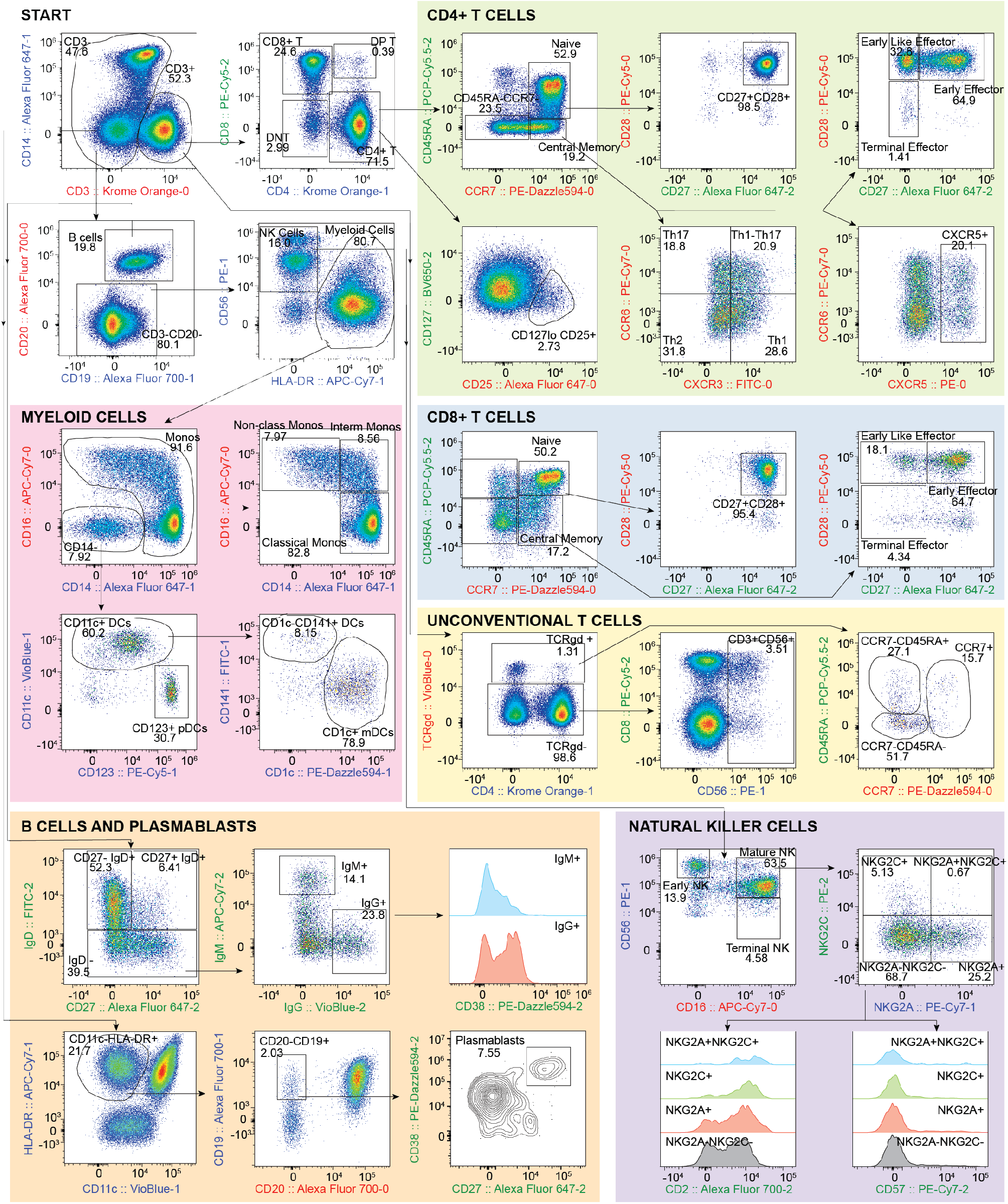
Manual gating of 33-marker, 3 cycle data. After excluding doublets and dead cells, populations of CD4+ and CD8+ T cells, unconventional T cells, myeloid cells, B cells, plasmablasts, and natural killer cells were identified via manual gating. Each major cell type was further differentiated into unique subsets using markers characteristically expressed on cells from healthy human populations. Axes labels are color coded for the cycle in which the marker was measured: Cycle 0 (red), Cycle 1 (blue), Cycle 2 (green).

## Discussion

Over the past decade, the prevailing approach to improve high-marker analysis with flow cytometry has been to add more excitation lasers and detectors to the instrument while developing newer fluorophores with dissimilar properties in optical absorption and/or emission. Current high-color commercial instruments (e.g., BD FACSymphony A5 and Cytek Aurora) use as many as 10 excitation lasers and 30 to 188 detectors to discriminate more than 30 different fluorophores at a time. However, this approach comes with significant cost and compromise. The spectral widths of organic fluorophores are typically 40 to 100 nm, and the detectable visible spectrum ranges from 400 nm to 800 nm. As a result, it is relatively routine to resolve up to about 10-15 different fluorophores, but beyond that, the assay difficulty increases nonlinearly for every fluorophore to be detected due to increasing SS (Supplementary Note 1). Recently introduced spectral detection can help distinguish similar colors, but spectral unmixing cannot compensate for photon shot noise and still leaves data spread^39,40^. In all, while an experienced flow cytometry user can design and optimize a 10-color experiment in approximately 1-2 weeks, a typical 30-color experiment requires several months to develop from scratch^8,41,42^. The time, technical expertise, reagent limitations, and cost needed for current high-marker panel designs have prevented many users from increasing the number of markers that are routinely measured.

Multi-pass flow cytometry based on optical barcoding alleviates this major bottleneck to high-marker analysis. First, it simplifies a highly complex panel into multiple, easier measurements, enabling more markers to be measured with fewer colors. This reduces the time and expertise needed to optimize a high-color panel and increases the margin of error afforded for the average user to acquire high quality data. Second, when no more than 10-15 easily distinguishable fluorophores are used in each measurement, there is no need for expensive, custom antibody reagents conjugated with exotic fluorophores of limited availability. Widely available and well-validated antibodies with common fluorophores can be used exclusively, significantly cutting reagent costs, lead times, and additional experiments needed for antibody validation. In addition, fluorophores which are the most widely available (e.g., PE, APC and FITC) can be re-used multiple times. Third, the reduced number of color channels simplifies the instrument design, instrument cost and operational complexity. Finally, by overcoming the fundamental limitation of SS when measuring many colors at once, multi-pass cytometry can exceed the maximum number of markers measured on a flow cytometer (currently, about 40 markers^34^), accelerating immunology and immune-oncology research by enabling analysis of more cell types at once.

Splitting panels across three cycles can drastically improve data quality. A conventional 30 color experiment requires monitoring of 30 detection channels at a time and optimizing 30 × (30-1) = 870 spillover matrix elements for compensation. Increased spillover inevitably causes increased SS, and high-parameter panels must be carefully designed to mitigate spreading error between co-expressing antigens. In comparison, a 3-cycle x 10 color experiment requires monitoring of 10 detection channels at a time and optimization of 3 × 10 × (10-1) = 270 spillover matrix elements, with most of these elements having relatively low spillover since less colors are used at a time. This dramatic overall reduction of spillover and corresponding spread contributes to major improvements of data quality. At the same time, investigators can also design panels such that co-expressing markers are split between cycles, thereby eliminating spread between these antigens altogether. When panels are split and acquired over several cycles, even markers conjugated to the same fluorophore multiple times will not require compensating and exhibit no spillover spread between them. In addition, splitting panels between cycles allows for re-using key fluorophores. For example, PE is a very bright and widely available fluorophore which could be used multiple times to detect low-antigen density markers.

Measuring tagged cells over multiple cycles inevitably introduces some reduction to cell yield. The cell tagging with PEI-coated LPs showed about 50% yield for live human PBMCs with 1 h of mixing (Fig. 2). This yield stems from the relatively lower internalization efficiency of certain cell types, such as T-cells. While minor differences in tagging efficiency for different cell types can be corrected in post-hoc analysis, overall cell yield may be a limitation for precious, heterogeneous cell samples such as primary patient samples. One approach to increase the tagging yield is to use antibody-based coupling of LPs to the cell surface^43,44^. This approach is similar to cell hashing employed for single-cell sequencing which uses antibodies targeting broadly and highly expressed markers. Our initial data shows that this can increase tagging yield to over 75% (Supplementary Fig. 1). Another area of potential improvement is matching yield, which in this study was around 70% per cycle. This yield is currently limited by the presence of free LPs that contributes to matching uncertainty, cells that are excluded because they are part of cell-cell doublets in at least one cycle, and LPs that may not be detected and/or become dislodged from the cell (see Methods). Optimization of the matching algorithm to include scatter and fluorescence data is also likely to improve matching yield.

Total analysis time is increased with the cyclic workflow due to the additional steps of LP tagging, cell capture, photobleaching and re-staining, apart from intentional time delay between measurements in time-lapse workflows. Currently, about 1 h is required for each additional cycle; however, only 10 minutes of this is hands-on time, and use of automated liquid handlers can shorten the current cell-processing time between cycles. LP tagging time and photobleaching time could be reduced further by using antibody-targeting and spectrum-optimized LEDs, respectively. Of note, LP-tagged samples can be fixed, stored, and measured the next day or whenever necessary, which may be a preferred workflow for panels with intracellular markers that require overnight fixation. Fixed samples can also be stored for batch analysis to reduce variability between specimens in large-scale clinical trials^45^. Re-interrogation of samples could also be useful for users who wish to use the results of a first cycle of measurement to inform the panel design of a subsequent measurement. The multi-pass workflow also requires that cells are run through the instrument multiple times, which imparts additional stress to the cells. However, marker expression should be minimally affected at the relatively low pressure (<3 psi) and low energy dissipation rate (~10^5^ W/m^3^) of our flow system^46–48^, and particularly sensitive markers can be deliberately acquired in the initial cycle.

The multi-pass workflow can also be leveraged in assays that require protocols or treatments which compromise fluorophore integrity. Methanol-based fixation often used to measure the phosphorylation state of intracellular proteins can quench protein-based fluorophores, rendering them unusable if staining precedes fixation^49^. Permeabilization buffers used to access intranuclear transcription factors often destroy signals from GFP and other fluorescent proteins^50^. In each of these cases, phenotyping cells in an initial cycle followed by fixation/permeabilization and subsequent measurement of intracellular markers enables measurement of all desired parameters without any compromise in signal or data quality.

Cellular barcoding and the multi-pass workflow also expand the utility of flow cytometry beyond static profiling to dynamic, time-resolved analysis of cells at high throughput. The ability to track and measure cells over time enables study of single-cell responses to stimulation, drug treatments, or other interventions. With conventional flow cytometry, only population differences can be identified when comparing control and treated samples. With time-resolved flow cytometry, the downregulation or upregulation of key biomarkers on individual cells can be identified and also quantified. The degree of change in expression of a particular biomarker on a particular cell could be especially useful for precision medicine applications. Tracking cells over multiple generations also enables the study of how protein expression changes as each cell divides or differentiates, with applications in tumorigenesis and stem-cell biology. We also anticipate that further LP barcoding innovation will enable us to couple flow cytometry with other optical instruments such as a fluorescence microscope^51^. The upstream or downstream integration of spatial and functional information of single cells, as illustrated in Fig. 1, promises to extend the ability to analyze single cells far beyond the current scope of flow cytometry.

## Acknowledgements

This work was supported in part by grants from the National Institutes of Health (R44-GM139504, R43-GM140527, DP1-EB024242, and R01-EB033155).

## Financial Interest Disclosure

All authors have financial interests in LASE Innovation Inc., a company focused on commercializing technologies based on optical barcodes. The financial interests of N.M. and S.H.Y. were reviewed and are managed by Massachusetts General Brigham in accordance with their conflict-of-interest policies.

## Author Contributions

S.J.J.K., S.F., M.D.F., S.C. designed the study. S.F., S.C., S.H.L, performed flow cytometry experiments. G.A. and S.H.L. performed laser particle tagging experiments. H.Z., N.H.M, and A.S.V designed and built the flow cytometer. H.Z. and N.H.M. designed and built the photobleacher. S.J.J.K., S.F., and M.D.F. analyzed and interpreted data. H.Z. and N.Ma. developed the matching algorithm. S.J.J.K. and S.H.Y. prepared the manuscript with input from all authors.

## Supplementary Figures

**Supplementary Fig. 1.**
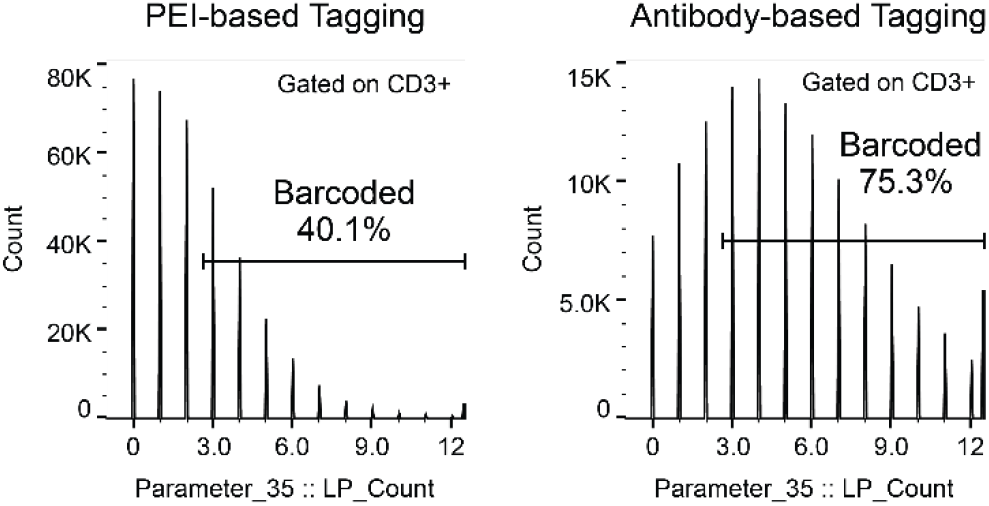
Comparison between LP tagging using a cationic polymer (PEI) and antibodies (beta-2-microglobulin).

**Supplementary Fig. 2.**
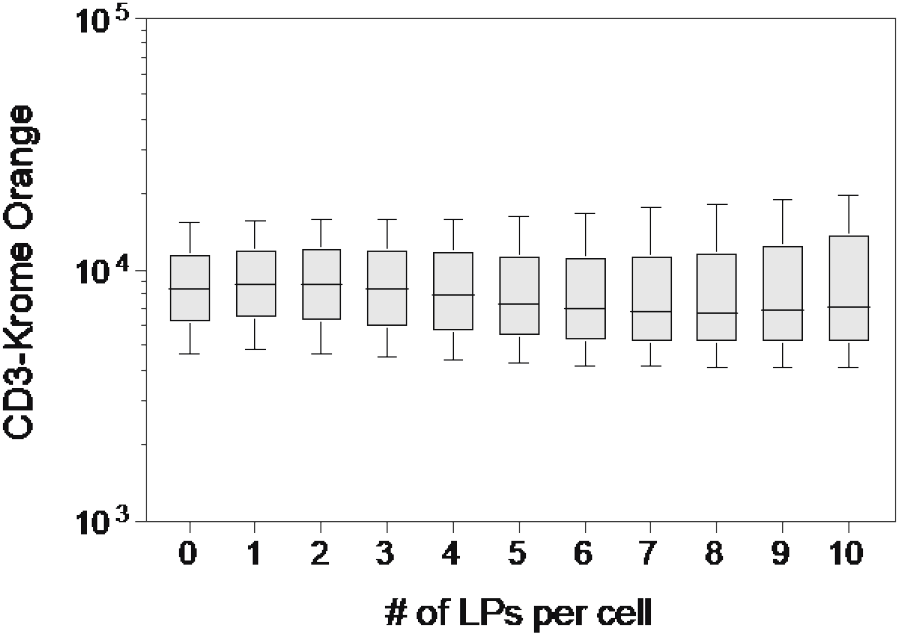
Dependence of CD3-Krome Orange fluorescence intensity on number of LPs per cell. Box plots show line at median, with error bars spanning the 10-90 percentiles.

**Supplementary Fig. 3.**
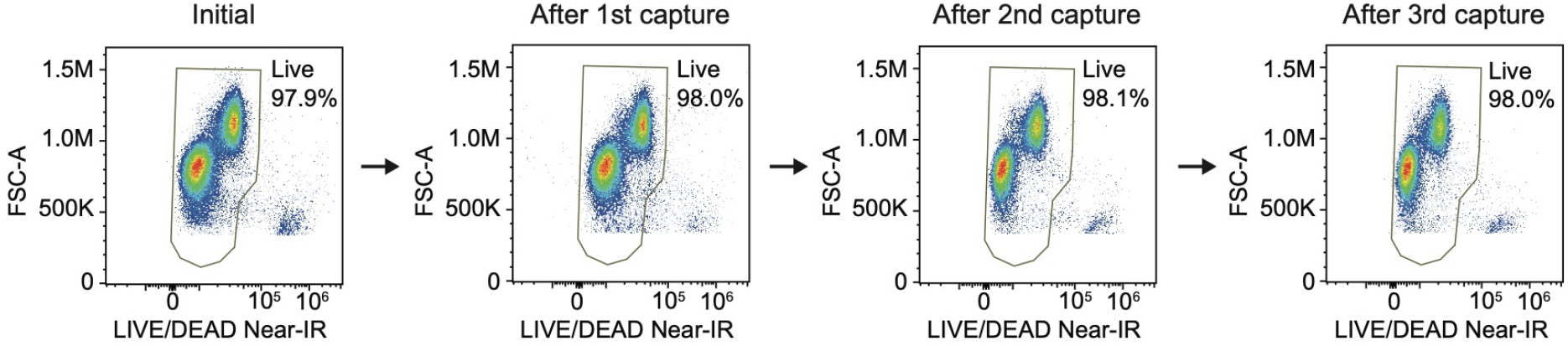
Cell viability of human PBMCs measured after 0,1,2, and 3 successive captures. Plots depict all CD45+ singlet events.

**Supplementary Fig. 4.**
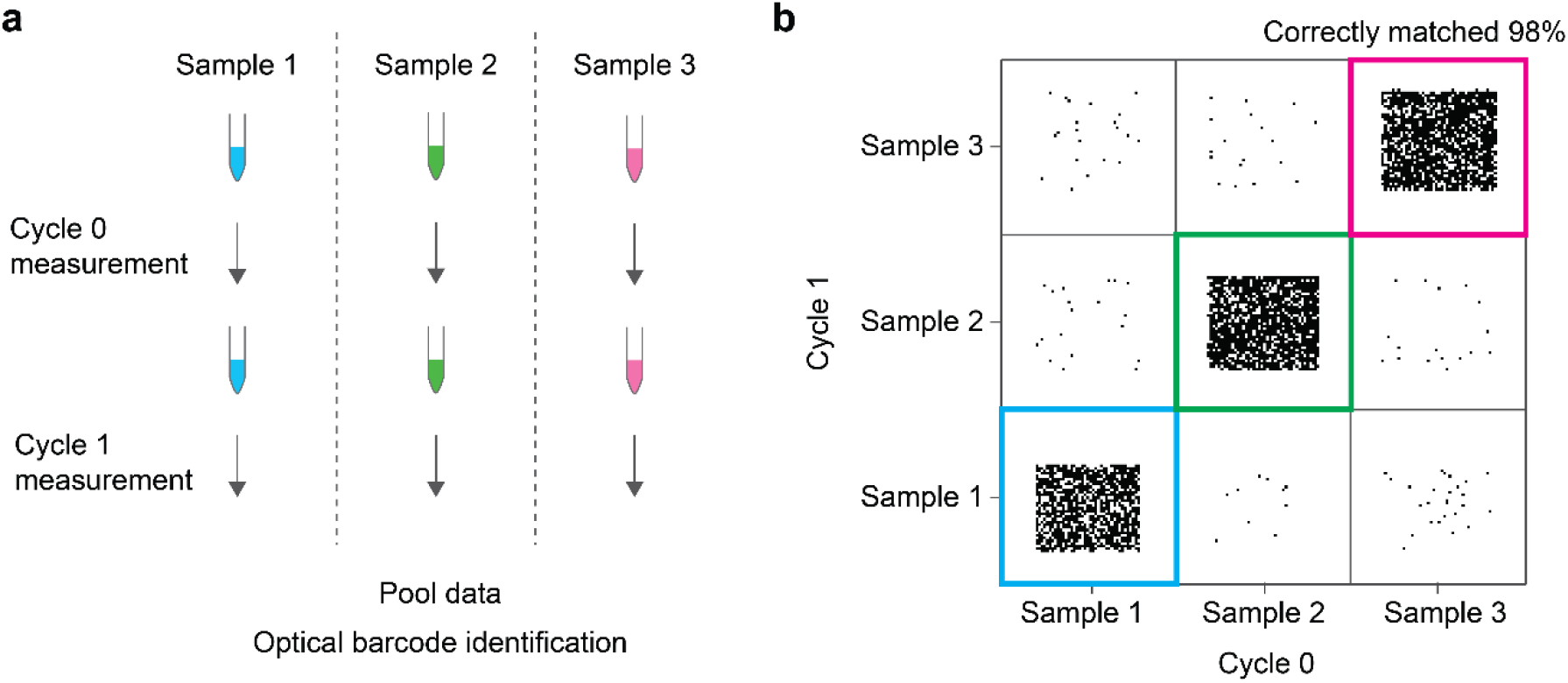
Validation of LP barcode matching. (a) Three cell samples each with ~200,000 barcoded cells were acquired and collected separately over 2 cycles. Data from the 3 samples were concatenated and matched to assess the accuracy of matching. (b) 98% of the matched cells were correct in maintaining sample identity across cycles. Plot only displays 6,000 cells out of ~360,000 for clarity.

**Supplementary Fig. 5.**
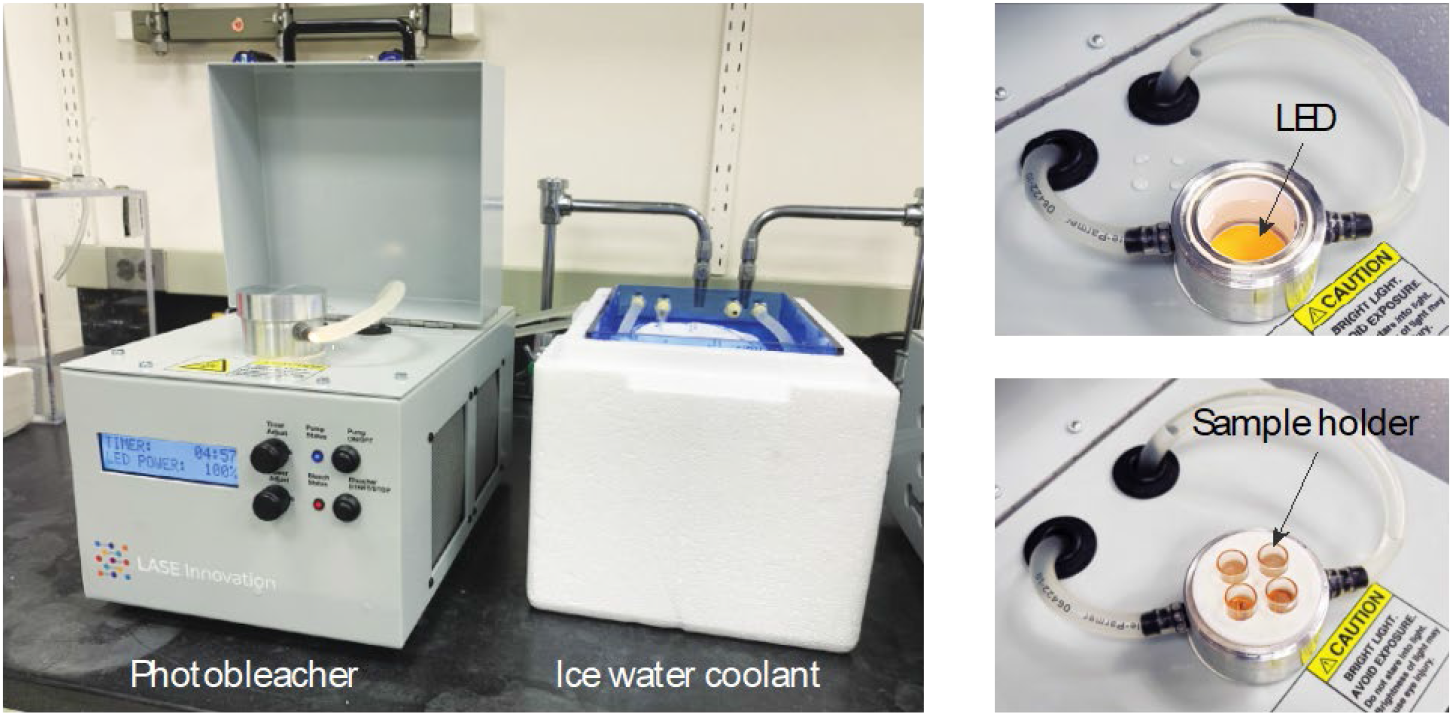
Image of custom-built photobleaching device in which to 4 samples are photobleached using a bright LED while being cooled to close to 4°C.

**Supplementary Fig. 6.**
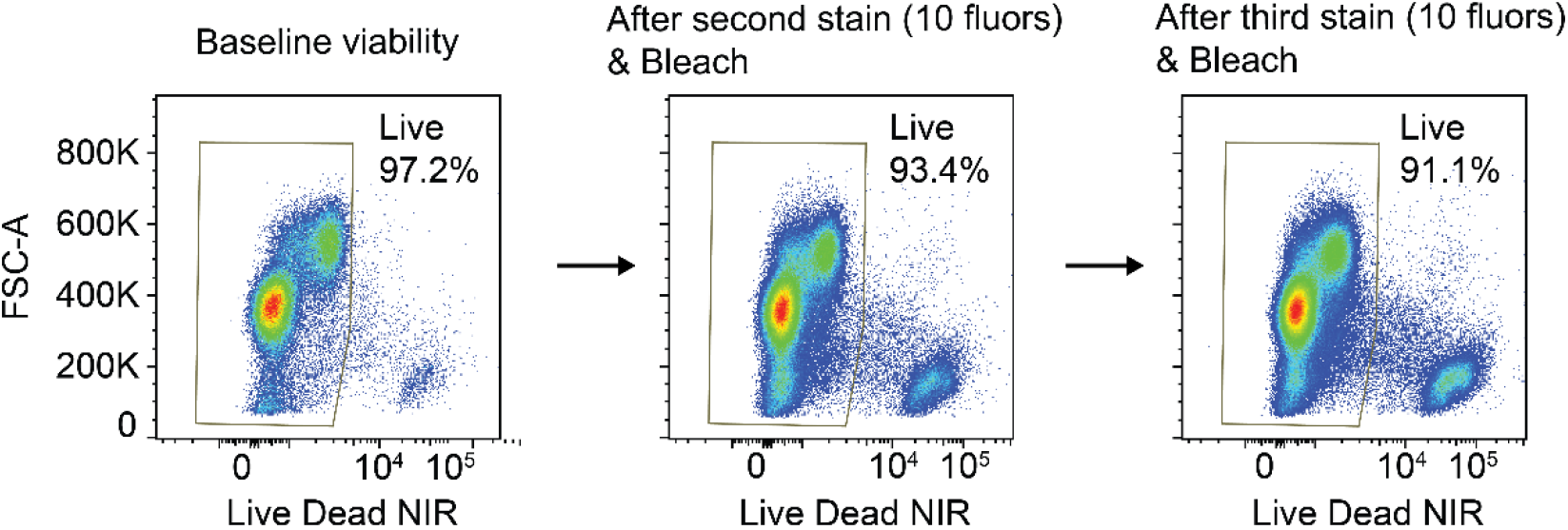
Viability of live human PBMCs following complete photobleaching of samples stained with 10 fluorophores (Cycle C0 in Table 1) and subsequently another 10 fluorophores (Cycle C1). The cell viability decreases around 3-4% per cycle. Plots depict all singlet events.

**Supplementary Fig. 7.**
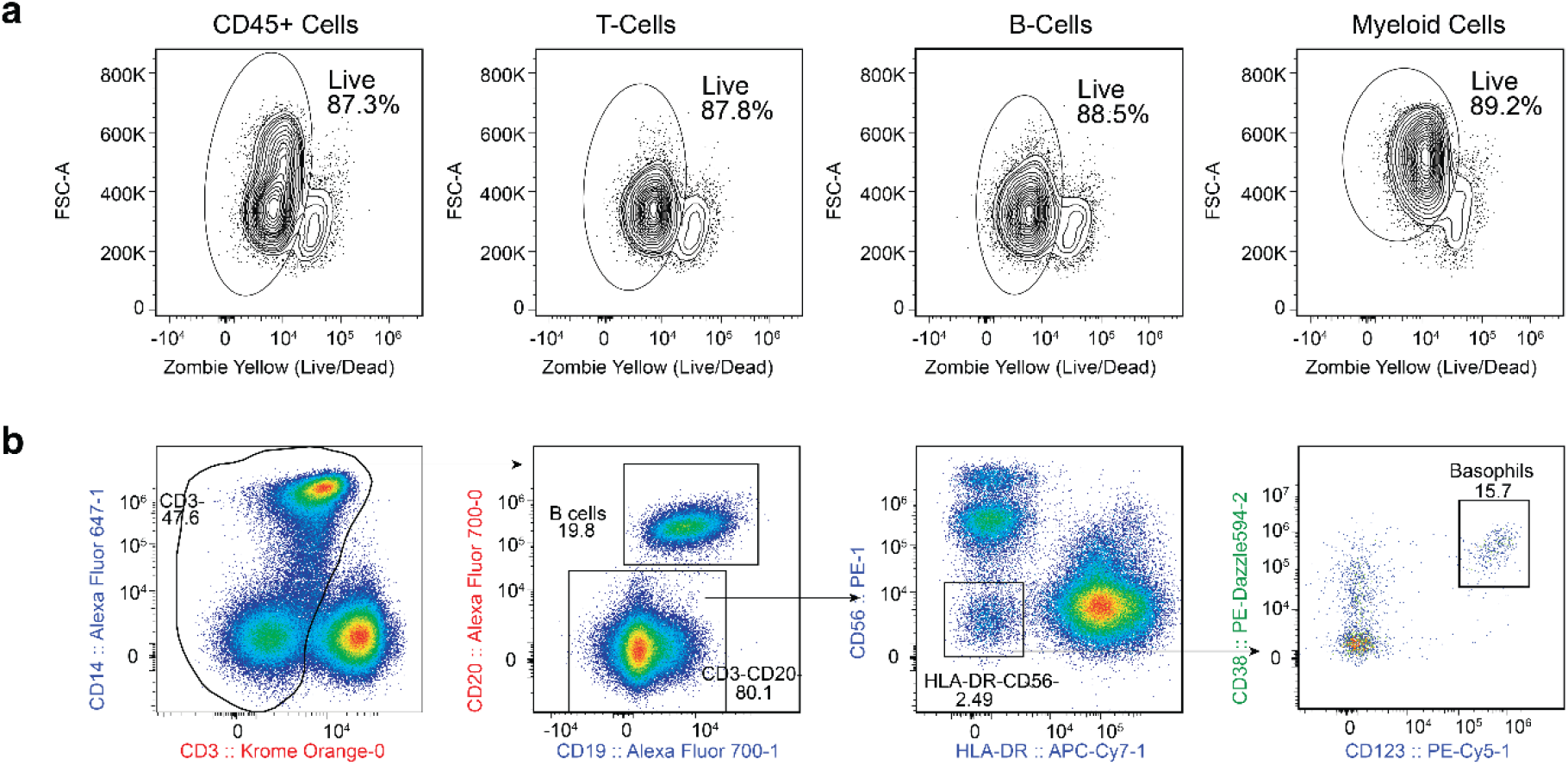
(a) Viability of cells measured in cycle 3 of the 33-marker, 3 cycle panel. Viability was measured using a Live/Dead dye for CD45+ cells, CD3+ T cells, CD20+ B cells and CD14+ Myeloid Cells. (b) Gating strategy for basophils.

**Supplementary Fig 8.**
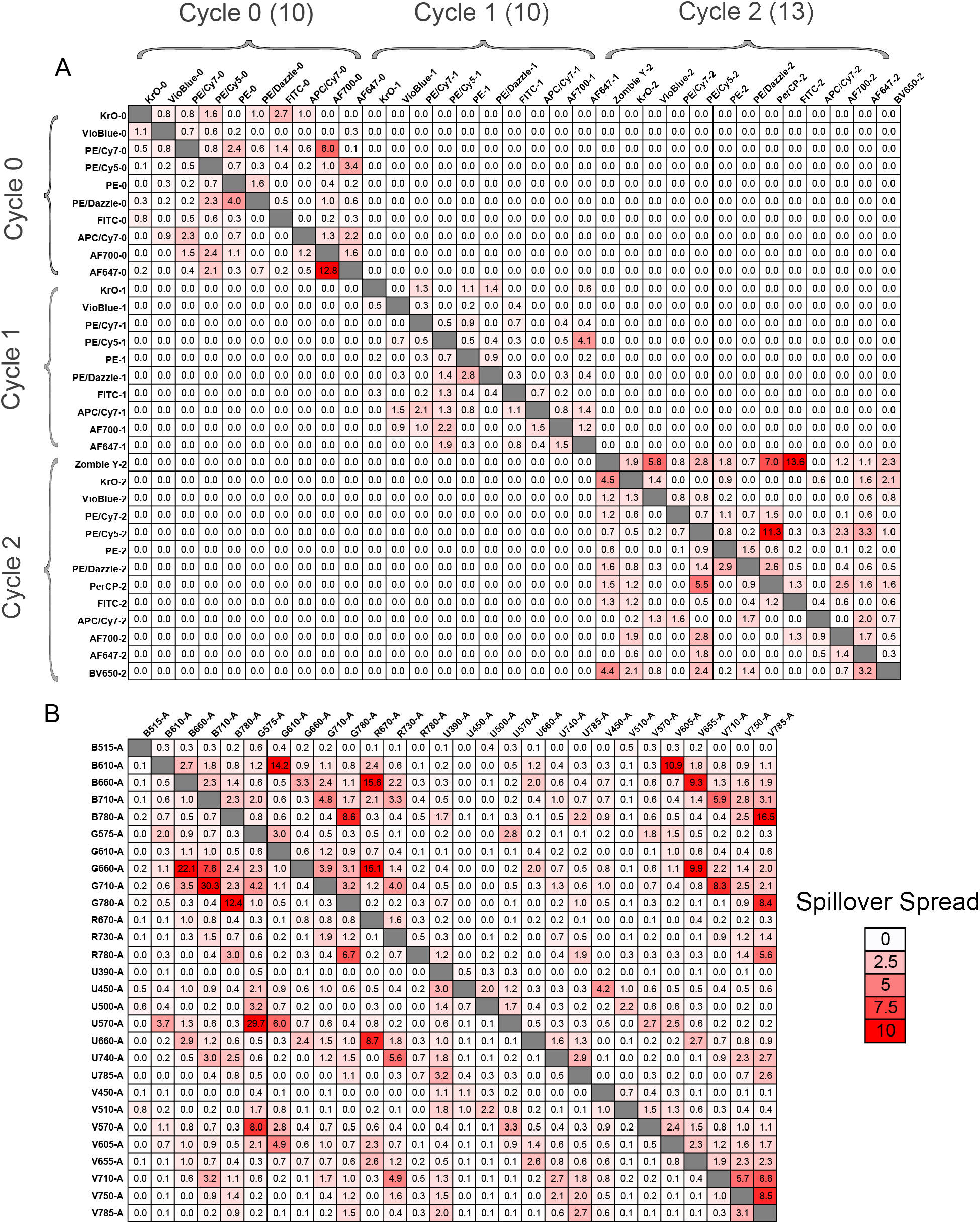
The spillover spread matrix (SSM) shows the extent of spread between fluorophores that occurs when emitted light from each fluorophore spills into non-primary detectors. (a) SSM computed for the 33-marker, 3 cycle panel demonstrated in this study.No spillover or spread occurs between fluorophores used in different cycles. (b) SSM computed for a published 28-marker, conventional flow cytometry panel (ref 23).

