## Supplementary Note 1 for "Laser particle barcoding for multi-pass high-dimensional flow cytometry"

**Supplementary Note 1: The total spillover spread in multi-color cytometry**

When a new fluorophore is added to a panel, its fluorescence spectrum has overlap with existing fluorophores, and as a result, increases the total spillover spread (SS). The magnitude of SS, $y$, as a function of the number of fluorophores, $x$, can be expressed as:

$$\Delta y(x) = y(x) - y(x-1) = \sum_{i=1}^{x-1}(\Gamma_{ix} + \Gamma_{xi}) \qquad (1)$$

where $\Gamma_{ix}$ and $\Gamma_{xi}$ are the elements in the rows and columns of the SS matrix containing the $x$-th fluorophore. $\Gamma_{ij}$ is approximately proportional, but not identical, to the spectral overlap between $i$-th and $j$-th fluorophores, but the precise relation is unimportant in the analysis here.

*A. Random choice of fluorophores*

Let us assume that all fluorophores have identical optical properties (absorption and emission linewidths) other than their center emission frequencies. When such fluorophores are added one by one randomly, we may express the above equation in terms of expectation values, $\langle \quad \rangle$ as:

$$\langle \Delta y(x) \rangle \equiv \frac{dy}{dx} \approx 2(x-1)\,\langle \Gamma_{ij} \rangle \qquad (2)$$

where $\langle \Gamma_{ij} \rangle$ represents the mean value of the matrix element. (The factor of 2 comes from the symmetry of the SS matrix, a sum along the row and the column containing the $x$-th fluorophore.) Eq. (2) gives

$$y = \langle \Gamma_{ij} \rangle (x-1)^2 \qquad (3)$$

$\langle \Gamma_{ij} \rangle$ is roughly proportional to the ratio of the total spectral range of detection to the linewidth of fluorophores. The quadratic dependence is obvious since the number of coefficients in the SS matrix grows in 2 dimensions, where each matrix element is $\langle \Gamma_{ij} \rangle$ in this random-addition case.

*B. Optimal choice of fluorophores*

In practice, fluorophores do not have the same linewidths, and the mean values of $\langle \Gamma_{ix} \rangle$ and $\langle \Gamma_{xi} \rangle$ vary depending on fluorophores. Fluorophores with broader emission spectra tend to have higher mean values of $\langle \Gamma_{ix} \rangle$ and $\langle \Gamma_{xi} \rangle$ compared to fluorophores with narrower emission spectra. In this more realistic case, an experienced panel designer does not use random fluorophores but considers spectral overlap as one of the critical metrics. For a given number of markers, the user usually ends up choosing fluorophores that result in minimal SS.

In this minimal-SS condition, from Eq. (1) we can write

$$\frac{dy}{dx} = 2(x-1)\,\Gamma(x) \qquad (4)$$

where $\Gamma(x) = (\langle \Gamma_{ix} \rangle + \langle \Gamma_{xi} \rangle)/2$ represents the mean overlap integral of the $x$-th fluorophore with the pre-populated $x-1$ fluorophores. More precisely, $\Gamma(x)$ corresponds to the sum of the matrix elements containing the $x$-th fluorophore (note that $\Gamma_{xx} = 0$). Under the minimal-SS protocol, $\Gamma(x)$ is a monotonously increasing function of $x$. (In the earlier random-choice case, $\Gamma(x)$ was a constant $\langle \Gamma_{ij} \rangle$.)

We have obtained the SS matrix from a set of 28 fluorophores used in a published T-cell panel (OMIP-060 and OMIP-068). From the matrix, we calculated $\frac{dy}{dx}$ and $y$ using the minimal-SS protocol as a function of $x$ from 1 to 28. Figure A shows the simulation result. The plot of $y$ appears in Fig. 6(e) in the main paper.

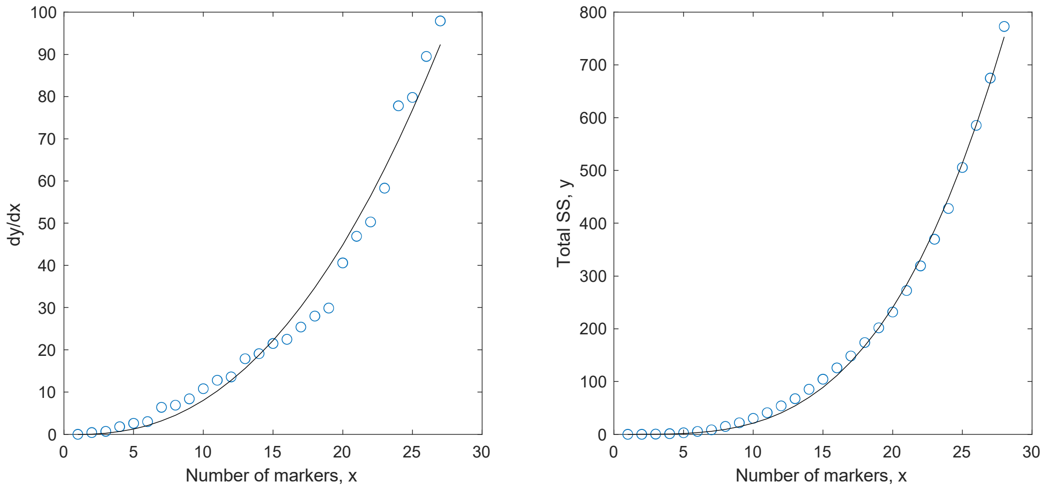


Figure A: Simulation result of the OMIP-068 panel.

The plot of $\frac{dy}{dx}$ is fit well with a single power-law function $A(x-1)^{2.3}$, which yields

$$f_\Gamma(x) \approx \frac{A}{2}\,(x-1)^{1.3} \qquad (5)$$

This shows that the increase of SS by adding $x$-th fluorophore increases with $x$. This is likely due to use of non-ideal fluorophores in high-marker panels. In practice, this is driven by limited antibody availability, need to use tandem fluorophores with broader emission and absorption linewidths and instrument constraints. The increment of SS by a fluorophore may be referred to as the spillover cost of the fluorophore (analogous to the chemical potential of a molecule to the free energy of the system). For example, the spillover cost is 0, 10 $A$, 24.6 $A$, and 41.6 $A$ for the 1$^{st}$, 11$^{th}$, 21$^{st}$, and 31$^{st}$ fluorophores, respectively.

The solution of Eq. (4) is

$$y \approx B(x-1)^{3.3} \qquad (6)$$

where $B = \frac{A}{3.3}$. The curve fit to the simulation data is excellent.

We find that the power-law dependence, $y \approx B(x-1)^k$, describes the characteristics of high-marker panels quite well. Figure B below shows the analysis of several other published panels, OMIP-060, OMIP-064, OMIP-067, OMIP-069, and OMIP-084. All of the results are fit reasonably well with a single power law function with an exponent in range of 2.8 to 3.5. A detailed interpretation of this finding is beyond the scope of this document.

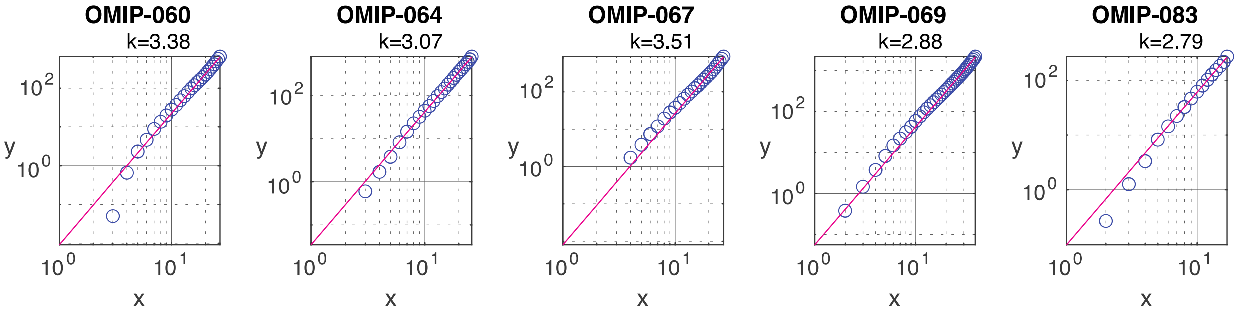


Figure B: Simulation results of various OMIP panels in the log scale. Lines, curve fit with $y = B(x-1)^k$; the best-fit $k$ values are indicated.

For comparison, we also calculated $y$ for the LASE's 3-cycle panels. The results are fitted with indices between 3 and 4, as shown below.

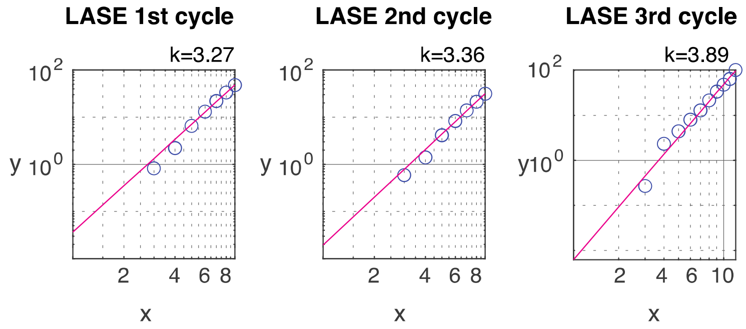


Figure C: Simulation results of the LASE 3-cycle panels in the log scale. the best-fit $k$ values are indicated.

*C. Cyclic measurement*

Now, consider $n$-time cyclic cytometry using only $x/n$ fluorophores. The total SSM at each cycle is $B\left(\frac{x}{n}-1\right)^{3.3}$. Since there are $n$ cycles, the total SS is given by

$$y_n = B\left(\frac{x}{n}-1\right)^k \times n \qquad (7)$$

For $\frac{x}{n} \gg 1,\ y_n \approx B\frac{(x-1)^k}{n^{(k-1)}}$. We find the ratio of the SS between the $n$-cyclic and non-cyclic cases to be

$$\frac{y_n}{y} \approx n^{-(k-1)} \qquad (8)$$

Let us consider the above 28-marker case with Where we use $k$ = 3.3. For $n = 3$ cycles, $\frac{y_n}{y} \approx 0.08$. This means that the SS in 3-cycle cytometry is 12.5 times lower than the SS in non-cyclic case, and this ratio is constant independent of the total number of markers.

Using $y(x) = y_n(x')$, we get $(x-1)^k = (x'-1)^k/n^{(k-1)}$ and find

$$x' = n^{(k-1)/k}\, x \qquad (9)$$

For $k$ = 3.3 and $n = 3$, we get $x' = 2.15\, x$. This means that using 3 cycles one can measure 2.15 times more markers with the same SS.

This simulation data of Eq. (7) obtained from the 28-marker panel appears in Fig. 6(e) in the main paper.

The number of fluorophores that can be used would be limited by the limited availability of fluorophores and undistinguishable spectral overlap with an existing fluorophore. Currently, the record experiment used $x = 40$ (OMIP-69, see Fig. B above). 3-cycle cytometry can extend this limit to 86. In principle, more cycles can further push the limit. With $n = 5$, up to 122 markers should be possible, requiring 24-25 fluorophores, with the same SS as the non-cyclic, 40-marker cytometry using 40 fluorophores.
